# Interacting networks of resistance, virulence and core machinery genes identified by genome-wide epistasis analysis

**DOI:** 10.1101/071696

**Authors:** Marcin J. Skwark, Nicholas J Croucher, Santeri Puranen, Claire Chewapreecha, Maiju Pesonen, Ying ying Xu, Paul Turner, Simon R. Harris, Julian Parkhill, Stephen D. Bentley, Erik Aurell, Jukka Corander

**Author notes:** these authors contributed equally (Marcin J. Skwark, Nicholas J Croucher). Corresponding authors: Jukka Corander; Erik Aurell.

## Abstract

Recent advances in the scale and diversity of population genomic datasets for bacteria now provide the potential for genome-wide patterns of co-evolution to be studied at the resolution of individual bases. The major human pathogen *Streptococcus pneumoniae* represents the first bacterial organism for which densely enough sampled population data became available for such an analysis. Here we describe a new statistical method, genomeDCA, which uses recent advances in computational structural biology to identify the polymorphic loci under the strongest co-evolutionary pressures. Genome data from over three thousand pneumococcal isolates identified 5,199 putative epistatic interactions between 1,936 sites. Over three-quarters of the links were between sites within the *pbp2x*, *pbp1a* and *pbp2b* genes, the sequences of which are critical in determining non-susceptibility to beta-lactam antibiotics. A network-based analysis found these genes were also coupled to that encoding dihydrofolate reductase, changes to which underlie trimethoprim resistance. Distinct from these resistance genes, a large network component of 384 protein coding sequences encompassed many genes critical in basic cellular functions, while another distinct component included genes associated with virulence. These results have the potential both to identify previously unsuspected protein-protein interactions, as well as genes making independent contributions to the same phenotype. This approach greatly enhances the future potential of epistasis analysis for systems biology, and can complement genome-wide association studies as a means of formulating hypotheses for experimental work.

**Author Summary:** Epistatic interactions between polymorphisms in DNA are recognized as important drivers of evolution in numerous organisms. Study of epistasis in bacteria has been hampered by the lack of both densely sampled population genomic data, suitable statistical models and powerful inference algorithms for extremely high-dimensional parameter spaces. We introduce the first model-based method for genome-wide epistasis analysis and use the largest available bacterial population genome data set on Streptococcus pneumoniae (the pneumococcus) to demonstrate its potential for biological discovery. Our approach reveals interacting networks of resistance, virulence and core machinery genes in the pneumococcus, which highlights putative candidates for novel drug targets. Our method significantly enhances the future potential of epistasis analysis for systems biology, and can complement genome-wide association studies as a means of formulating hypotheses for experimental work.

## Introduction

The study of co-evolution in recombining populations of bacteria has been limited by the scale and polymorphisms present in population samples for which whole genome sequences are available. Even the most recent population genomic studies of bacterial pathogens have been constrained in this respect, such as focusing on a particular genotype(1-3), biasing sampling towards particular clinical outcomes(4-6), or surveying organisms in which limited genetic diversity and strong linkage disequilibrium (LD) masks the signals of shared selection pressures(7, 8). For whole genome-scale modeling of co-evolution, sampling should span the entirety of diverse, recombining species in an unbiased manner.

The first organism satisfying all the above-mentioned desiderata is *Streptococcus pneumoniae* (the pneumococcus), for which over 3,000 genome sequences from a well-defined, limited study population were recently published(9). As the pneumococcus is an obligate nasopharyngeal commensal and pathogen, the bacterial population was evenly sampled through a structured survey of the hosts. The diverse multi-strain population structure, coupled with the naturally transformable nature of *S. pneumoniae*, results in low LD across the genome. Hence this set of pneumococci can be considered as an ideal set for detecting genes that evolve under shared selection pressure.

Analyzing sets of co-evolving polymorphisms is a powerful means of identifying sites that interact directly, through protein-protein contacts, and indirectly, through epistatic interactions that affect the same phenotype. The former type of selection pressure has previously been studied on the scale of individual proteins. It has been known for more than 20 years that the correlations of amino acids in two columns in a multiple sequence alignment (MSA), contain exploitable information and provide a non-trivial predictor of spatial proximity(10, 11). Detection of co-evolving mutations in genomes is in the statistical sense analogous to this structural prediction problem, as both phenomena are consequences of joint selection pressures. The latest advances in computational structural biology have shown that by changing the modeling framework from correlations to high-dimensional model learning one can improve protein contact predictions significantly(12-15). Furthermore, including considerations of epistatic interactions between sites has recently been shown to significantly improve the mapping between genotype and phenotype for a beta lactamase protein(16).

Co-evolving sites do not necessarily directly interact, however; instead, changes at distinct sites may represent selection for a particular phenotype determined by multiple polymorphic loci. However, the complexity of the possible set of interactions has mostly limited previous analyses of epistasis to viral datasets of limited diversity; nevertheless, these studies have shown epistasis to be an important factor in evolution. An application of a phylogenetically-informed method to influenza subtypes H1N1 and H3N2 identified patterns of substitutions associated with the emergence of resistance to oseltamivir(17), and many sites were found to be undergoing coordinated evolution within the hepatitis C virus(18). However, the non-linear increase in the number of interactions as the genome length and diversity rises limits the application of such methods to the study of bacterial populations. In recent work, pairwise statistical correlation analysis was demonstrated to successfully reveal certain types of co-evolutionary patterns across the genome for 51 *Vibrio parahaemolyticus* isolates(19). Nevertheless, pairwise analyses of association are subject to Simpson’s paradox which may cause spurious links to emerge(20-22), and furthermore, the necessity of correcting for a quadratically increasing number of multiple hypothesis tests seriously hampers the statistical power to detect the true positive associations as sample sizes increase.

Here we demonstrate a new method able to identify co-evolving polymorphisms from bacterial genome sequence alignments named genomeDCA and made freely available at www.helsinki.fi/bsg/software/genomeDCA. By considering the evolution of polymorphic sites simultaneously and using the inference tools for regularized statistical model learning one avoids both the problems that drive high levels of false positives and negatives when the number of pairwise interactions grows. The methods introduced here may offer a powerful alternative to traditional GWAS analyses for multiple unknown phenotypes in the emerging era of massive population sequencing for bacteria.

## Results

### Genome-wide identification of coupled loci

The whole genome alignment used in this analysis consists of short-read data for over 3,000 systematically-sampled pneumococcal isolates (9) each aligned to the reference sequence of *S. pneumoniae* ATCC 700669 (23). Filtering this alignment for biallelic loci at which the minor allele frequency was >1% and >85% of isolates had a base called identified 81,560 polymorphic sites. Of these, 88.3% were within protein coding sequences (CDSs), a slight enrichment relative to the 87.2% of the *S. pneumoniae* ATCC 700669 reference sequence that was annotated as CDSs. Following the genome-wide linkage analysis (see Methods), estimates of association strength were retained from 102,551 couplings (Table S1). A Gumbel distribution fitted to this sample (Fig. 1; μ = 0.096 and β = 0.028) significantly diverged from the empirical data above a coupling strength of 0.129. The 5,199 couplings (Table S2) exceeding this threshold were considered as putative epistatic interactions; these affected 1,936 sites, 89.0% of which were within CDSs. As closely proximal sites were excluded from this analysis, these coupled sites had a mean separation of 587.4 kb, with only two sites less than five kilobases apart (Fig. 2). Hence these associations are unlikely to be artifacts of genetic linkage.

Fig 1 Divergence between theoretical and empirical distributions of coupling strengths between sites. Left panel (A) shows the two distributions such that the vertical axis corresponds to the log10 probability of a coupling coefficient exceeding the value of the curve on the horizontal axis. The dashed vertical line depicts the significance threshold; 5199 out of 102,551 couplings exceed the threshold. Right panel (B) displays the absolute difference between the fitted cumulative Gumbel distribution and the empirical cumulative distribution (on log10-scale) as a function of the coupling strength. The dashed vertical line marks the smallest coupling (0.129) which has a difference of more than six standard deviations among the first 50,000 empirical-Gumbel differences.

Fig 2. The 5199 significant couplings shown by lines connecting genomic positions. The thickness of lines is proportional to the number of linked positions within the corresponding chromosomal elements. The red markers show the positions of sites identified in an earlier GWAS study of resistance determining variation in the pneumococcal genomes. The green markers indicate locations of protein coding sequences where significant couplings are present.

### Strong epistatic links between penicillin-binding proteins

The putative epistatic interactions were used to generate a network (Fig. 3). The nodes, each corresponding to a CDS, were colored according to function and scaled according to the number of epistatic links with which they were associated. The edges were weighted according to the number of interactions between CDS pairs. Most of the annotated functional categories were represented in the network, with the notable exception of mobile genetic element genes. This was almost entirely the consequence of the absence of informative sites in these regions of the genome, owing to their high variability across the population. By contrast, the functional category most over-represented in the dataset was surface-associated proteins. Although previous work has suggested immune selection might drive epistasis between antigens (24), in fact this enrichment was entirely the consequence of selection for antibiotic resistance. Of the 4,617 links represented in this network, 3,578 (77.5%) involved 175 sites found in one of three genes encoding penicillin-binding proteins (PBPs; Fig. 3A): SPN23F03080 (*pbp2x*), SPN23F03410 (*pbp1a*) and SPN23F16740 (*pbp2b*). These PBPs have been experimentally demonstrated to be the major determinants of resistance to beta lactams (25), with changes to each individual protein reducing its affinity for beta lactam antibiotics while retaining its the ability to bind its natural substrates(26, 27). This link between genotype and phenotype was also identified by a genome-wide association study (GWAS) using this dataset(28). Of the 858 sites found to be significantly associated with beta lactam resistance by this GWAS, 403 were within these three coding sequences for PBPs; of these, 216 met the criteria to be analysed in this study. Correspondingly, 161 of the 175 sites identified within the same genes by this analysis were significantly associated with beta lactam resistance by the GWAS (Fisher exact test, OR = 112.1, *p* < 2.2x10^−16^), indicating the couplings between sites in these genes represent the set of changes that distinguish pneumococci with differing sensitivities to beta lactam antibiotics. This was in congruence with the distribution of these alleles across the population (Fig S1), which also confirmed that none of the identified associations correlated with the expansion of a single clone, indicating the method’s correction for the effects of population structure was effective.

Fig 3 Network of coupled protein coding sequences. This undirected network shows all significant couplings between protein coding sequences (CDSs). Each node is a CDS, colored according to its functional annotation, and scaled according to the logarithm of the number of significant coupled loci it contained. Edges are weighted according to the logarithm of the number of significant coupled loci linking two CDSs. (A) Network component containing the genes *pbp2x*, *pbp1a* and *pbp2b*. (B) Network component containing the *smc* gene. (C) Network component containing the tRNA synthetase gene *pheS* and a coding sequence for another putative tRNA-binding protein. (D) Network component containing the genes for *pspA* and *divIVA*.

Changes in these genes are strongly epistatic, as for pneumococci to develop non-susceptibility to a broad range of beta lactam antibiotics, alterations of all three PBPs are necessary. Consequently, the emergence of multidrug-resistant pneumococci is associated with transformation events that alter all three over very short evolutionary timescales (29-31). Another factor that might underlie both the concerted changes at all three loci, rather than a gradual emergence of resistance, as well as the non-uniform distribution of coupled sites across these genes (Fig. 4) is the potential for alterations in only one protein disrupting direct protein-protein interactions:. As these proteins perform similar functions on the same substrate, and are all co-localised to the cell wall, it has been hypothesised that they function as constituents of a multi-enzyme complex(32). Some evidence from co-immunoprecipitation and crosslinking experimental work has supported this idea(33, 34). Hence the distribution of coupled sites between the PBPs was investigated in greater detail.

Fig 4 Distribution of couplings between sites in different PBPs. The red markers are defined as in Fig. 2.

### Structural distribution of coupled sites in PBPs

Couplings were identified between all three PBPs, although almost 95% involved *pbp2x*. This might reflect that *pbp2x* has to be altered for resistance to both penicillins and cephalosporins, whereas modifications of *pbp1a* are important primarily for cephalosporin resistance (35), and modifications to *pbp2b* are important primarily for penicillin resistance (36). Alternatively, if these proteins do interact, these data suggest *pbp2x* would be central to any potential protein-protein interactions. The coupled sites are distributed broadly across *pbp2x*, with the exception of the PBP dimerization domain, despite the identification of sites within this region by GWAS.

Co-evolving sites within *pbp1a* are more narrowly distributed and involve stronger interactions with *pbp2x* than with *pbp2b* (Fig 4, Table S2). The links between PBP1A and PBP2X are distributed between two domains and a further structural analysis (Fig 5,Table S3) showed how the identified positions in *pbp1a* with the strongest couplings were located in strand β-4 and the loop connecting strands β-3 and β-4, near the transpeptidase active site but not overlapping with the conserved catalytic residues. The counterpositions in *pbp2x* showed a spatially less focused pattern with linked positions near the active site, including the typically conserved active site residues Ser395 and Asn397, as well as links to positions not in direct spatial proximity of the active site, but structurally linked to that region through helical secondary structure elements. Alterations in the active site surroundings can interfere with inhibitor binding with only minor effect to catalytic function as was evident for substitutions in the identified β-3/β-4 loop region residues 574-577 which were strongly linked to beta-lactam resistance(37). Resistant strain *pbp1a* likewise showed amino acid substitutions at the identified positions 583 and 585 in strand β-4 in comparison to a beta-lactam susceptible strain(37) (resistant strain PDB code 2V2F). Position 580 at the N-terminal end of β-4 (typically a proline) in *pbp1a* was linked to position 363 in *pbp2x*, which is part of an ionic interaction (Glu363 to Arg372) present in all current crystal structures of *S. pneumoniae pbp2x* (see Methods for details) and may play a structural role by stabilising the α-2/α-4 loop region proximal to the *pbp2x* active site. Residues at PBP2X positions 401, 404, 412 and 413 are all buried within the protein, but are connected to active site residues Asn397 and Ser395 via helix α-5. It is possible that these positions are implicated in active-site shaping as well. For *pbp2b*, structural mapping of the top ranking co-evolved sites revealed two major groupings: positions in the α-2/α-4 loop region that, similarly to the *pbp1a* case, partially cover the active site, and positions that, similarly to the *pbp2x* side of *pbp2x* – *pbp1a* couplings, were spatially more distant but structurally linked to the active site. As in *pbp1a*, observed flexibility in the α-2/α-4 loop region proximal to the active site points to a potential role in antibiotic resistance of this structural feature(38).

Fig 5 Structural models of *pbp1a*, *pbp2x*, *pbp2b* with the 100 strongest couplings listed in Table S3 indicated. The figures show the transpeptidase domains of each PBP with catalytic/active site residues shown in cyan and coupled positions as sticks with other colors. Active site bound antibiotic/inhibitor is rendered as a space-filling volume when present in the crystal structure. Panels A-D depict: *pbp1a* with couplings to *pbp2x*, green colored residues are coupled with green residues in panel B; orange colored residues in B are coupled with both green and yellow residues in A (A), *pbp2x* with couplings to *pbp1a* (B), *pbp2x* with couplings to *pbp2b* in orange (C), *pbp2b* with couplings to *pbp2x* in orange (D).

### Associations with other resistance phenotypes

The PBPs are confined to a single network component that contains ten other proteins. Seven of these are found in close proximity to the three PBPs, and likely represent sequences altered when resistance-associated alleles of the PBP genes were acquired through transformation events, which often span tens of kilobases (29, 31). However, it is also possible these could play a role in ‘compensating’ for deleterious side-effects of the changes in the PBP proteins. One of these CDSs proximal to a penicillin-binding protein gene is *mraY*, directly downstream of *pbp2x* and encoding a phospho-N-acetylemuramoyl-pentapeptide-transferase also involved in cell wall biogenesis (Table S2, Fig 3). It was previously predicted that mutations in this transferase associated with beta lactam resistance could represent compensatory changes ameliorating the costs of evolving beta lactam resistance(28). Another CDS, *gpsB*, is shortly upstream of *pbp1a* and encodes a paralogue of DivIVA that also plays an important role in peptidoglycan metabolism (39).

The three proteins in the network component that were not proximal to a PBP-encoding CDS were *dyr* (also known as *folA* or *dhfR*), encoding dihydrofolate reductase, and three nearby genes (Table S2, Fig 3). Mutations in the *dyr* gene cause resistance to trimethoprim (40). The earlier GWAS study(28) found a significant association between both *dyr* and *folP* with beta lactam resistance, despite no functional link to such a phenotype, nor any likely reason why they would directly interact with PBPs. Hence the detected interaction between *dyr* and the *pbp* genes is most likely explained by the co-selection for resistances that have accumulated in the same genetic background, resulting in the multi-drug resistant genotypes observed to have emerged over recent decades(41).

### Couplings between core genome proteins

To identify other functional roles that might underlie the distinct sets of couplings represented in Fig. 3, a gene ontology (GO) analysis was performed for each network component containing more than two nodes. This identified five significant signals, including that for penicillin-binding associated with the previously described component (GO:0008658, Fisher’s exact test, OR = 337.3, *p* = 0.00048 after Benjamini-Hochberg correction). However, the strongest association was that of the largest network component, containing 384 CDSs (Fig. 3B), with ATP binding activity (GO: 0005524, Fisher’s exact test, OR = 2.89, *p* = 8.75x10^−7^). Other GO terms significantly associated with this component were GO:0005737, corresponding to cytosolic localisation (Fisher exact test, OR = 3.44, *p* = 0.00010 after Benjamini-Hochberg correction), and GO:0016021, corresponding to integral membrane proteins (Fisher exact test, OR = 2.32, *p* = 0.043 after Benjamini-Hochberg correction). These associations partly reflect the preponderance of cytosolic ATP-hydrolysing tRNA synthetases, of which enzymes for the processing of eleven amino acids were present among these CDSs, and membrane-associated ATP-hydrolysing ABC transporters. The most highly connected node in the component, linking to 22 other CDSs, was another ATPase. SPN23F11420, encoded the Smc protein, is critical in organizing the chromosome and forms the basis of a multi-protein complex in both prokaryotes and eukaryotes (42). Hence this large diverse set of coupled CDSs included many components of the essential cytosolic machinery, the interactions of which are critical to the basic functioning of the cell.

The fourth significant enrichment of GO terms also involved tRNAs (GO:0000049 - tRNA binding; Fisher exact test, OR = 869.1, *p* = 0.00048 after Benjamini-Hochberg correction), which applied to a component containing three nodes (Fig 3C). One corresponded to *pheS*, a phenylalanyl tRNA synthetase, while the other was SPN23F19340, annotated as encoding a tRNA binding protein of unknown function. Attempting to identify a more specific functional prediction using the CDD database (43) we found this protein possessed a “tRNA_bind_bactPheRS” domain, specifically involved in processing phenylalanyl-tRNAs, and only otherwise found in PheT, which directly interacts with PheS in the phenylalanyl-tRNA synthetase. Hence this coupling may represent a previously unexpected direct protein-protein interaction.

The CDS directly downstream of *pheS*, encoding a putative membrane-associated nuclease (SPN23F05260), was coupled to a tRNA methyltransferase adjacent to *pspA* in a separate network component (Fig. 3D). The *pspA* gene, encoding a surface-associated protein involved in pathogenesis and immune evasion, was itself present in the same network component, and engaged in some of the strongest coupling interactions in the dataset. These linked to the *divIVA,* encoding a cell morphogenesis regulator paralogous with GpsB (39), and three CDSs upstream of *ply*, encoding the major pneumococcal toxin pneumolysin(44) which, like *pspA*, is critically important in pneumococcal virulence and upregulated during infection(45). These three CDSs (SPN23F19480-SPN23F19500) are likely to play a role in localising or transporting *ply* from the cytosol into the cell wall(44). This could be the consequence of these virulence proteins engaging in interactions at the surface of the cell.

This set of interactions was also detectable when a bipartite network was constructed that displayed couplings between coding sequences and upstream untranslated regions (Fig S2). In general, these components mirrored those in Fig. 3, suggesting the same couplings were being represented, rather than epistatic interactions involving direct protein-DNA interactions. Correspondingly, neither DNA binding (GO: 0003677) nor RNA binding (GO:0003723) were enriched in this network.

## Discussion

Natural selection continuously performs experiments in bacterial populations, leading to purging of deleterious sequence variation and maintenance of beneficial mutations. Laboratory experiments provide the gold-standard method for establishing underlying mechanisms among observed variable sites. However, they also necessitate the definition of a measurable phenotype, which may be a daunting task for many complex traits relevant for survival and proliferation of bacterial strains. The exponentially increasing size of the genome sequence databases provide a valuable resource for generation of hypotheses for experimental work. In eukaryotes, GWAS methods have been used for more than a decade to probe DNA variation which could explain phenotypic differences(46, 47). In bacteria, use of GWAS for this purpose is of much more recent origin and has been demonstrated to hold a considerable promise in the light of more densely sampled populations(28, 48-50). However, GWAS is not the only way in which wealth of bacterial sequence information has been proposed to be used to gauge which genes could potentially be targets of positive selection and to generate hypotheses for experimental work. For example, Li et al. screened genome sequences of closely related pairs of isolates in a densely sampled pneumococcal population which would differ at particular genes of interest to provide candidate targets for phenotypic tests(51).

By leveraging from the most recent advances in computational protein structure prediction and statistical machine learning, we have been able to introduce a method that promises to complement the popular GWAS approach for understanding how polymorphisms affect phenotypic variation. This work identified many different coupled sites across the genome, which network analysis revealed to define separate clusters of genes involved in resistance, virulence, and core cell functions. Our study represents the first attempt to use statistical modeling to fully exploit large-scale bacterial population genomics to identify patterns of co-evolution in sequence variation. Importantly, as we have demonstrated, it is not necessary to specify the relevant phenotypes *a priori* for this approach to work successfully. Hence this a single analysis with this method may simultaneously reveal co-selected sites for a multitude of different traits, and provide an indication of their relative influence on evolutionary patterns, as illustrated by the results on the pneumococcus. Since our approach is widely applicable to data generated by bacterial population studies, it has considerable potential to identify important targets for experimental work to gain system level understanding about the evolution and function of bacteria.

## Materials and Methods

### Genome data

The 3,156 genomes used in our study are obtained from the study by Chewapreecha et al(9). We used their genome alignment of the total length 2,221,305 bp, in which 388,755 SNP loci were present, out of which 134,037 loci had a minor allele frequency (MAF) of at least 0.01. Out of these we selected 81,560 loci that had coverage of at least 84.1%, i.e. not more than 500 genomes with a gap/unresolved base pair at the considered sequence position. As the analysis focused solely on the biallelic loci, the observed nucleotides have been replaced, so that the entire alignment was composed of only three letters (two representing observed allele (major/minor) and one gap/unobserved allele). This was done in the interest of reducing the number of parameters of learnt models approximately 3-fold.

### Regularized model learning

The model we use for joint evolution of SNP loci is the Potts generalization of the Ising model(52), the latter characterizing interactions between Boolean variables based on an exponential family of distributions with parameters for every possible pair of loci. In the three state case (minor/major allele and gap) the Potts model specifies couplings between pairs of loci through 3x3 parameter matrices, and consequently the parametric dimensionality grows by the number of loci squared, i.e. is here on the order of ~10^10^, whereas the number of observations is on the order of 10^3^. Models are therefore regularized as discussed in Ekeberg et al(53). The relevant predictions from the model are the strongest interactions obtained and a threshold for significant interactions was determined using the statistical theory of extreme value distributions as explained below.

### Pseudolikelihood inference and parameter scoring

Pseudolikelihood was originally introduced in the early 1970’s to enable estimation of parameters in spatial statistical models with intractable likelihood functions(54, 55). This inference technique has experienced a strong revival in the recent years for high-dimensional applications where the number of possible model parameters greatly exceeds the number of observations, known as the ‘small n, large p’ problem(56). In particular, pseudolikelihood provides consistent estimators of the model parameters unlike the traditional variational inference methods(57). The outcome of the inference is a set of real numbers *Jij* describing the interactions between loci *i* and *j*. We score these numbers by their absolute values |*J_ij_*|, which corresponds to the Frobenius norm scoring of interaction matrices in the contact prediction problem. The pseudolikelihood method allows an efficient correction for population structure by the reweighting scheme used for MSA in protein analysis(53), which ensures that highly similar sequences are not artificially inflating the support for direct dependence between alleles.

### Resampling procedure

The number of parameters in our model is far larger than has hereto been considered in analogous studies. Also, neighboring loci are most often in LD which has a confounding biological effect on their interaction. For both these reasons, we have chosen to dissect the pneumococcal core chromosome into approximately 1,500 non-overlapping segments and apply pseudolikelihood inference on subsets of the loci, chosen randomly from each genomic window of average size of 1500 nt. For each such sample we learned about 10^6^ parameters, scored them as described above and saved only the 3000 largest interaction parameters to reduce memory consumption. The whole re-sampling and model fitting procedure was repeated 38,000 times to ensure stable inference about the parameters. Interaction estimates were averaged for any pairs of sites that occurred multiple times among the saved parameters, which resulted in 102,551 pairs of sites with non-negligible coupling coefficients from the aggregated re-sampling results (Table S1). This set is approximately five orders of magnitude smaller than the set of all possible interactions for the 81,560 considered loci. Software implementing both the resampling procedure and the parameter inference is made freely available at www.helsinki.fi/bsg/software/genomeDCA.

### Choice of significance threshold for interactions

To select a list of highest scoring interactions among the 102,551 estimates which are unlikely a result of neutral and sampling variation in the studied population, we employed the statistical theory of extreme value distributions (58). Since in each resampling step the largest 3000 parameters were saved, these can under a null model of random interactions between loci be considered as a sample from an extreme value distribution, such as the Gumbel distribution. We fitted a Gumbel distribution to the distribution of the estimated parameters using least squares minimization between the fitted distribution and the empirical rank distribution of the coefficients located between 25% and 75% quantiles. Fig 1A shows the fitted distribution which has a remarkably good fit to the vast majority (95%) of the coefficients. To select a threshold for a significant deviation from the null model we identified the first value for which the predicted curve was more than six standard deviations (SD) away from the empirical distributions (Fig 1B). The SD was estimated using the deviances for the 50,000 smallest coefficients. The 5199 couplings exceeding the threshold 0.129 are listed in Table S2 and in addition the 500 strongest couplings in Table S3. The latter were used in the structural plots for the PBPs.

A resampling-based analysis of haplotypes generated randomly from a population by merging alleles sampled from the marginal allele frequency distribution of each SNP locus showed that couplings as large as those exceeding the threshold chosen for the coefficients in the original data were never encountered (Fig S3). In the analysis we used 5000 replicates of the haplotype re-sampling based on the same chromosomal windows as in the analysis of the original data. Hence, our approach was concluded to maintain a strict control of false positive interactions for unlinked loci stemming from population sampling variation.

### Functional analysis and structural modeling

Networks were displayed and analysed using Cytoscape (59). GO terms were inferred from applying Interpro scan (60) and CD-search (43) to the *S. pneumonia* ATCC 700669 genome [EMBL accession: FM211187]. These were matched to network components, and a Fisher exact test used to test for enrichment of 139 instances of GO terms that featured in a network component twice or more, relative to the CDSs that contained sites analyzed in this study, but not found to include a significantly coupled loci. The *p* values were corrected for multiple testing using the method of Benjamini and Hochberg (61).

Crystal structures of *S. pneumoniae* PBPs with the following IDs: 2C5W (*pbp1a*), 2WAF (*pbp2b*), 2ZC3 (*pbp2x*) were retrieved from the Protein Data Bank(62) (www.rcsb.org; accession date January 8, 2016) and visualized in The PyMOL Molecular Graphics System, Version 1.8 Schrödinger, LLC. Inferred co-evolving sites were visualized using the Circos software(63).

## Acknowledgements

We thank Professor Brian Spratt for helpful comments on the manuscript.

## Author contributions

E.A. and J.C. designed the study, M.J.S, N.J.C., S.P., C.C., M.P., Y.X., E.A., J.C. analyzed data, P.T., S.R.H., J.P., S.D.B. provided biological expertise and interpretation of the results, M.J.S, N.J.C., S.P., J.P., E.A., J.C. wrote the manuscript, all authors approved the final manuscript.

## Supplementary information captions

**Figure S1.**
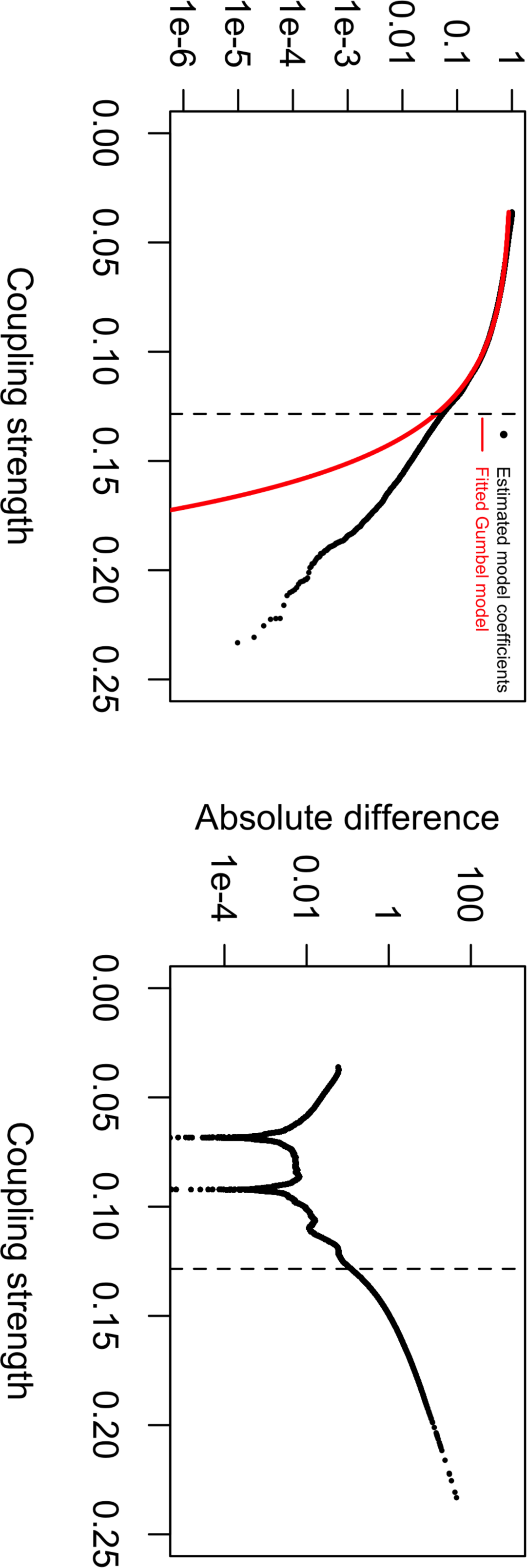
Comparing the distributions of coupled sites in penicillin-binding protein genes with those identified by a genome-wide association study for polymorphisms associated with beta lactam resistance. (A) Domain annotation of the PBP2X, PBP1A and PBP2B proteins, based on analysis with the Pfam database. (B) Distribution of coupled loci identified by this analysis. The columns corresponding to polymorphic loci identified as being significantly coupled with others by this analysis are coloured according to the base present in each isolate in the collection, ordered according to the whole genome phylogeny shown on the left. (C) Distribution of loci found to be significantly associated with beta lactam resistance through a genome-wide association study of this Maela population by Chewapreecha *et al*. Only those sites that match the inclusion criteria for this study (i.e. biallelic with <15% of sites missing across the population), and are within the three displayed genes, are shown.

**Figure S2.**
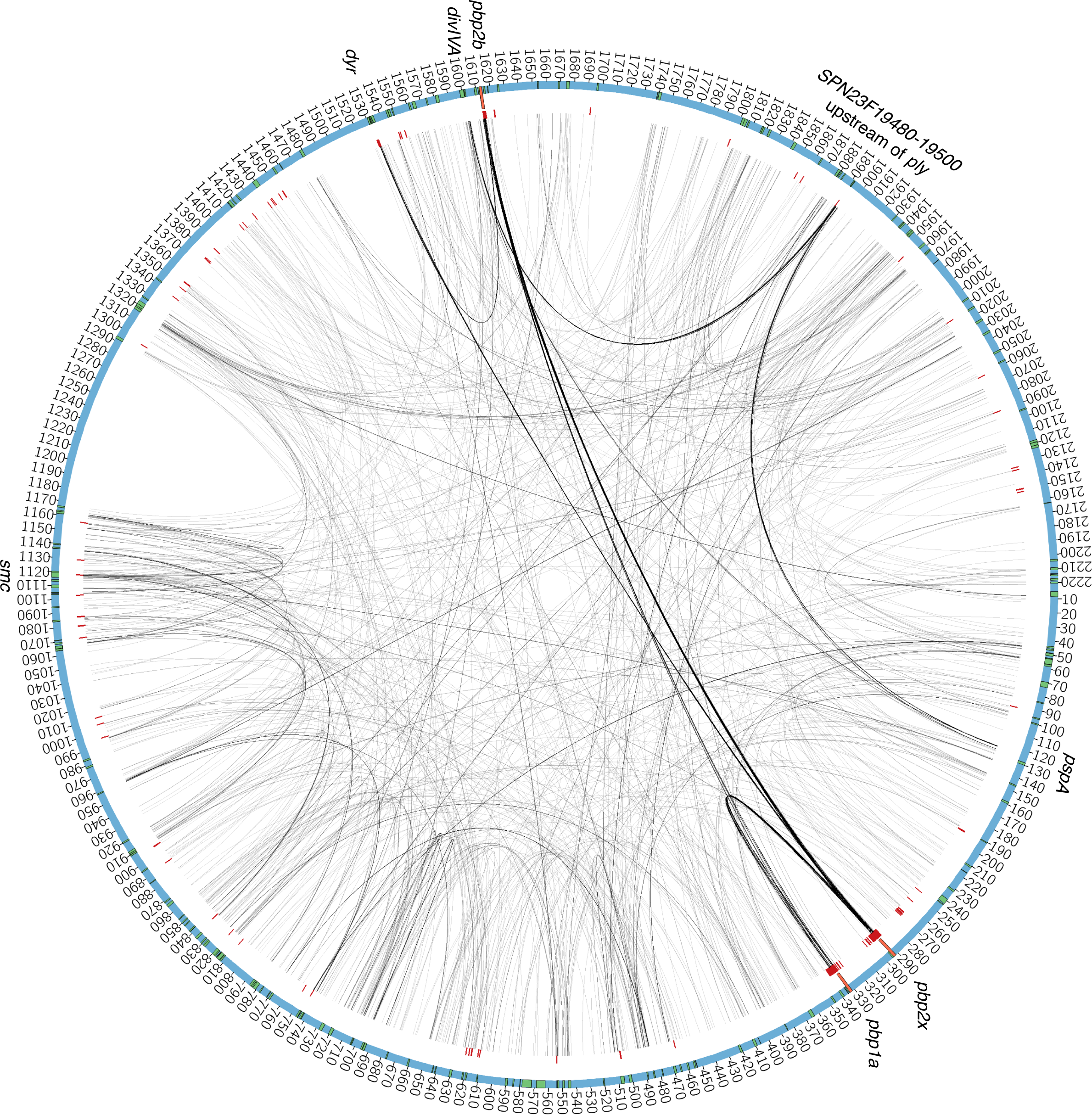
Bipartite network showing the couplings between coding sequences and upstream untranslated regions. The network is displayed as in Fig. 3, except that untranslated regions are shown as black triangles. (A) Network component showing the links between *pbp2x*, *pbp2b* and the upstream regions around *pbp1a*. (B) Network component showing the coupling between *smc* and upstream regions. (C) Network component showing the *pspA* and *divIVA* genes.

**Figure S3.**
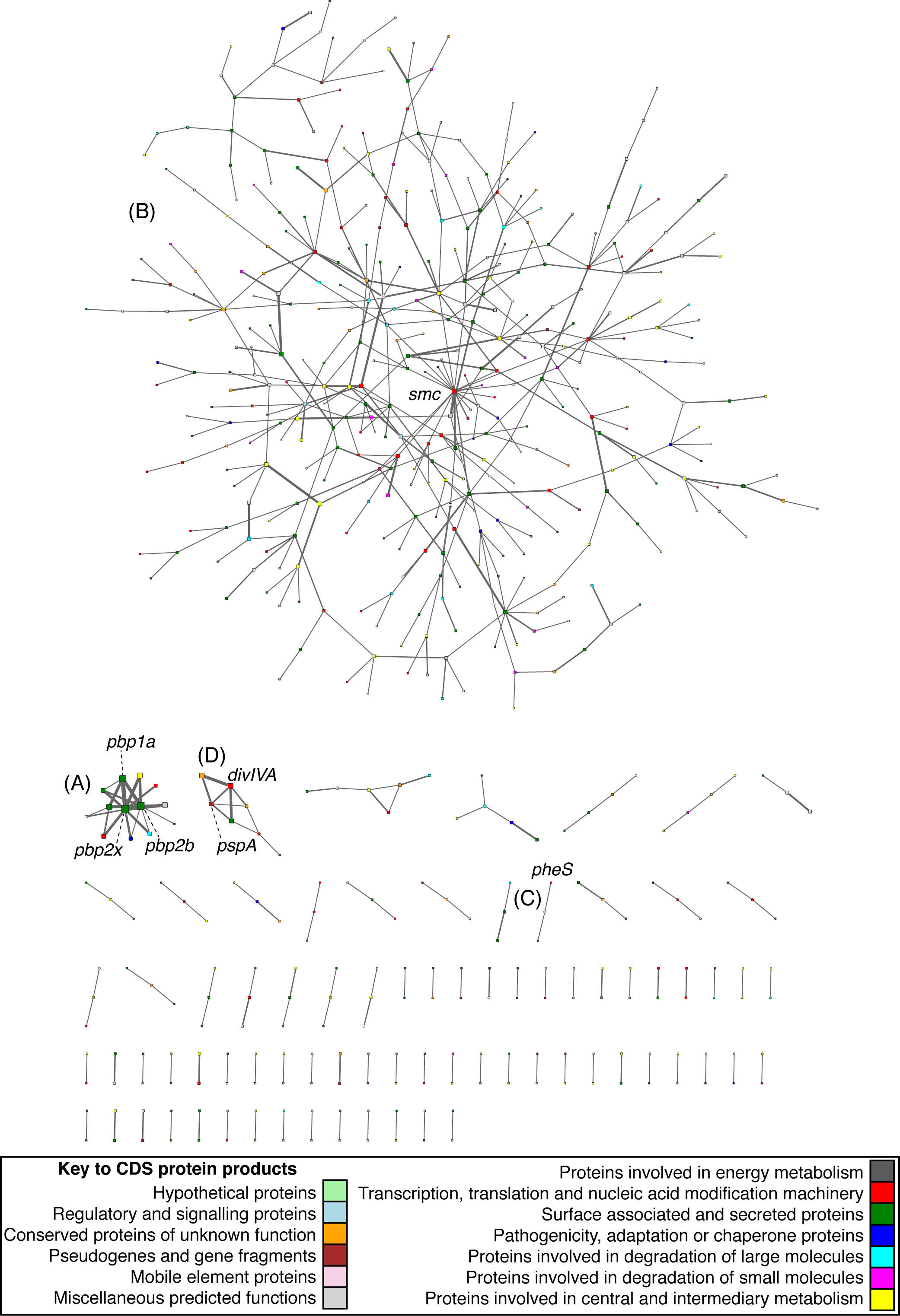
Distributions of estimated coupling coefficients for the original data and for haplotypes generated randomly from a population by merging alleles sampled from the marginal allele frequency distribution of each SNP locus. The red curve is generated from 5000 replicates of the haplotype re-sampling based on the same chromosomal windows as used in the analysis of the original data.

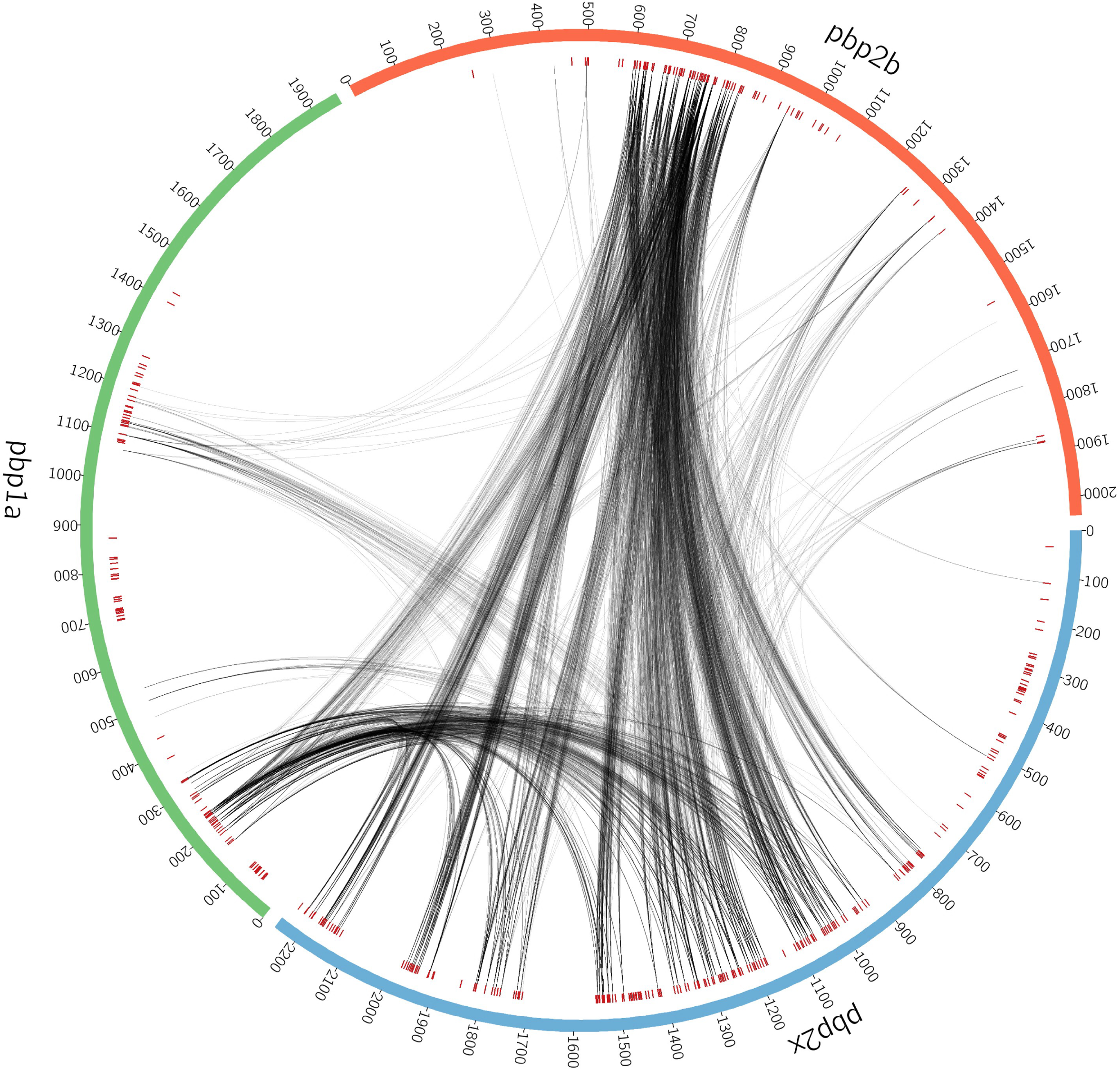

